# Contrastive Learning Improves Generalization on M2OR Olfactory Binding Dataset

**DOI:** 10.64898/2026.11.04.763508

**Authors:** Logan Snider, Oliver Nakano-Baker, Devin Mackenzie

## Abstract

Olfaction-based biosensors stand to have profound impacts in fields such as medical diagnostics, national security, and drug discovery. A promising mode of vapor phase biodetection mimics volatile organic compound-olfactory receptor (VOC-OR) interactions with target-specific protein or peptide probes; however, lack of experimental coverage remains a core bottleneck. Current computational approaches aim to improve prediction of VOC-OR binding to reduce experimental burden, and current machine learning approaches fail to demonstrate a level of generalization that addresses current engineering challenges. Here, using the Molecular to Olfactory Dataset (M2OR) and Insect Olfactory Binding Protein Database (iOBPdb) datasets, data partitioning methods and their performance are evaluated on a neural network and contrastive learning framework on such VOC-OR binding datasets, varying: model training, architecture, and structure of the input data distribution. It was found that holding out certain chemical or proteomic motifs from the training data diminished model performance with a two layer neural network model. The model was able to generalize to state of the art performance (0.88 AUCROC) on every data split by applying a contrastive learning objective defined by shared molecular constituents and a stochastic sampler. This suggests the contrastive objective provides a strong inductive bias for neural networks to predict novel VOC-OR pairs. These findings highlight that model performance on pairwise biological interaction datasets can be inflated through identity leakage from constituent overlap, and emphasize the need for more robust evaluation protocols. Furthermore, we show contrastive regularization mitigates dataset degeneracy through enforcement of compatibility in the models latent geometry that may extend to other binder target-datasets where identity leakage confounds evaluation.

## 1 Introduction

Biological olfactory systems transcend current engineering capabilities via specialized olfactory receptor (OR) proteins that selectively bind to volatile organic compounds (VOCs), leading to the transduction and processing of a signal [1]. Terrestrial animals also use olfactory binding proteins (OBPs) to bind to ligands for selective transport to sensory neurons. [2] Together, ORs and OBPs form a synergistic chemosensing architecture where OBPs selectively modulate VOC diffusion while ORs recognize and transduce via neural action potential. [3] Current engineered sensors often rely on narrow and static functional responses, while biological systems operate under dynamic conditions with redundancies and adaptations that enable sensitive and flexible functional response. These properties can enable robust sensing in noisy and sparse chemical environments. To replicate these capabilities, efforts must be taken to reconcile the differences between the stochastic and adaptive nature of biological systems and the rigid architectures presented in current engineering approaches.

We currently lack molecular structural information of VOC-OR systems. Transmembrane proteins like ORs are notoriously difficult to structurally characterize. Novel methods like CryoEM have been able to do so in some cases, but operate at a very low throughput compared to the immense OBP, OR and VOC interaction space and are not a general solution for scalable screening and selection of ORs. [4] AlphaFold 3 predicts ligand binding conformations; however, in general, machine learning methods are limited by the quality and coverage of the training data distribution given to train the model. [5] The problem remains that trans-membrane proteins, thus the VOC-OR binding domain, have been shown to be under sampled in the training data for these models, hence current structural prediction underperforms for VOC-OR binding while remaining relatively robust for other classes of proteins.

Without reliable structures, VOC–OR models must learn from functional and binding assays. Datasets like M2OR pool results from multiple wet-lab studies, as well as most VOC–OR binding datasets, this heterogeneity can degrade model performance and reproducibility. [6] Models may also learn statistical shortcuts rather than biologically meaningful features. [7]

Robust modeling should take effort to harmonize evaluation with the data’s structure and desired application. Binding data take three forms, quantitative affinity, predicted structure, and binary classification, each reflecting distinct biochemistry. Every affinity datum pairs a binder (usually a protein) with a ligand; though the binder–ligand space is sparse, individual proteins and ligands recur throughout each dataset. Random pairwise splits in iOBPdb, M2OR, and similar datasets thus place shared binders and ligands in both train and test sets, letting models memorize binding tendencies instead of learning the underlying chemistry and biology. Because predicting novel binders and ligands is often the whole point of these models, splits must instead hold out specific chemical motifs, protein classes, or binder–ligand pairs, each defining a fundamentally different learning task.

As modern machine learning has developed, the role of representation learning and regularization techniques has become increasingly pertinent to help models mitigate dataset bias. VOC-OR machine learning is a space where heterogeneous data is the norm, underscoring the importance of understanding the biases and interactions between the data collection, modality and parsing of data, as well as model architecture and regularization, as each aspect can significantly affect performance.

In this work we investigate how to carefully curate models to generalize to samples outside of the training data distribution on iOBPdb and M2OR, with methods transferrable to other ligand-binder datasets.

## 2 Background and Related Works

To predict a binding affinity between a protein and VOC, each chemical species must first be encoded as a vector. A classic algorithmic encoding method for chemicals is Morgan fingerprints, which enumerate sub-graphs of the molecular structure in a sparse 1 dimensional vector. [8] Algorithmic protein encoding methods include OneHot or categorical encodings, where vectors reference amino acids via positional or integer encoding, respectively. [9] Kernel methods have been success-fully applied to protein property prediction, but they are mostly limited to interpolation of input data distributions, often introducing considerable complexity to leverage effectively in deep learning pipelines. [10]

Transformer encoders have proven to be powerful molecular encoders, including ChemBERTa and ProtBERT used in this work. [11, 12]. In recent years, transformers have become popular as language models that encode sequences or strings (e.g., the Simplified Molecular Input Line Entry System [SMILES] for molecules, amino acid sequences for proteins) as dense vector representations for downstream machine learning tasks. Other deep learning methods like Graph Neural Networks and Recurrent Neural Networks remain widely used. [13, 14] Transformers can be powerful transfer learning layers, but they inherit bias from their training data, and may suffer diminished performance when used to predict samples outside of their original training data distribution. [15] Algorithmic methods do not inherit bias, but have less representation power than deep learning methods. The best encoder method varies with task and dataset size. [16]

Another primary consideration in ML VOC-OR binding prediction is the choice of how to best utilize encoded features in downstream tasks. Primitively, the data may be concatenated and passed into a classifier like a neural network or tree-based model. Attention-based techniques may be used to extract relevant motifs from encoded data which are then passed to the classifier model. In addition, semantic definitions can be used to construct graph representations of molecules and proteins and machine learning can be applied to these graphs for structurally informed machine learning.

Contrastive learning is a self-supervised learning method in which multiple augmented views of a sample are created and labeled as “positive pairs” and all other samples are labeled as “negative pairs”. An auxiliary contrastive learning objective is utilized to enforce clustering of positive and separation of negative pairs in the model’s latent representations. By applying these transformations, the model learns latent features that are invariant to various semantics-preserving transformations. Another benefit is that the effective number of data points in a dataset is scaled by the number of augmentations that one provides per sample, providing a way to generate more data points in data sparse regimes. Contrastive learning can be applied in a self-supervised manner, where the model is trained to maintain this self-consistency of the latent space before introducing labels. [17] Contrastive learning may also be applied in a supervised manner by using the task and the contrastive objective simultaneously; the model learns self-consistent geometry as it learns to predict the targets [18].

## 3 Methods

This work evaluates the influence of data encoding and dataset stratification on machine learning classification of olfactory protein binding affinities to VOCs on the M2OR and iOBPdb datasets. This was done using four encoders applied in pairs, three data stratification strategies, a downstream two layer neural network, and with both traditional and contrastive loss functions.

M2OR is an olfactory binding dataset compiled from 45 literature sources of ORs and VOCs. It contains 771 unique molecules and 1,402 unique protein sequences, with a total dataset size of 53,444 VOC-OR pairs, each classified as binding or non-binding. M2OR defines VOCs by their SMILES string and ORs by their protein sequence. [19]

iOBPdb is another olfactory binding dataset compiled from 181 literature sources of olfactory binding proteins (OBPs) and VOCs. It contains 622 unique molecules, and 382 unique protein sequences, totaling 8001 pairs spanning 91 insect species. iOBPdb defines VOCs by their SMILES string and OBPs by their protein sequence, with a reported dissociation constant (Kd) [20]. This was converted into a classification problem by designating all samples with a Kd below 10 as positive binders.

To represent VOCs and ORs from their semantic definitions, molecular and proteomic features were generated for downstream model training using either a pretrained transformer or algorithmic methods. These encoding methods were applied separately to the VOC and the OR and then concatenated before being passed into a downstream classifier. Bidirectional Encoder Representations from Transformers (BERT) architectures ChemBERTa and ProtBERT were chosen for their proven robust performance across bio- and cheminformatics tasks [21, 22]. ChemBERTa and ProtBERT are both Masked Language Models that were trained to capture contextual dependencies to predict tokens (amino acids for proteins, and atoms or bonds for molecules) that are left out of the input. Their weights were frozen, and the SMILES and sequences were passed into the frozen models to obtain the encoded representations for VOCs and ORs respectively. Algorithmic methods were 1024-bit Morgan fingerprints for VOCs and integer (categorical) encoding for ORs. Morgan finger-prints encode molecular graph substructures into a fixed-length binary bit vector, and an integer encoder encodes each amino acid in the sequence as an integer. An integer encoder was used after it outperformed one-hot encoding in early explorations of the M2OR dataset.

Three splitting strategies tested the ability of models to generalize outside of their training distribution. The “random split” was implemented where points were assigned to train, test, and validation sets randomly without replacement. “Stratified” splits selected data points by their unique VOC or OR identity, assigning train, test, and validation sets by randomly selecting unique VOC or OR identifiers and collecting all corresponding samples into one set. The stratified splits are intended to assess model generalization on each side of the protein-VOC pair, with protein-stratified data testing novel proteins and VOC-stratified data testing novel VOCs. Figure 1 illustrates the three split strategies.

**Figure 1:**
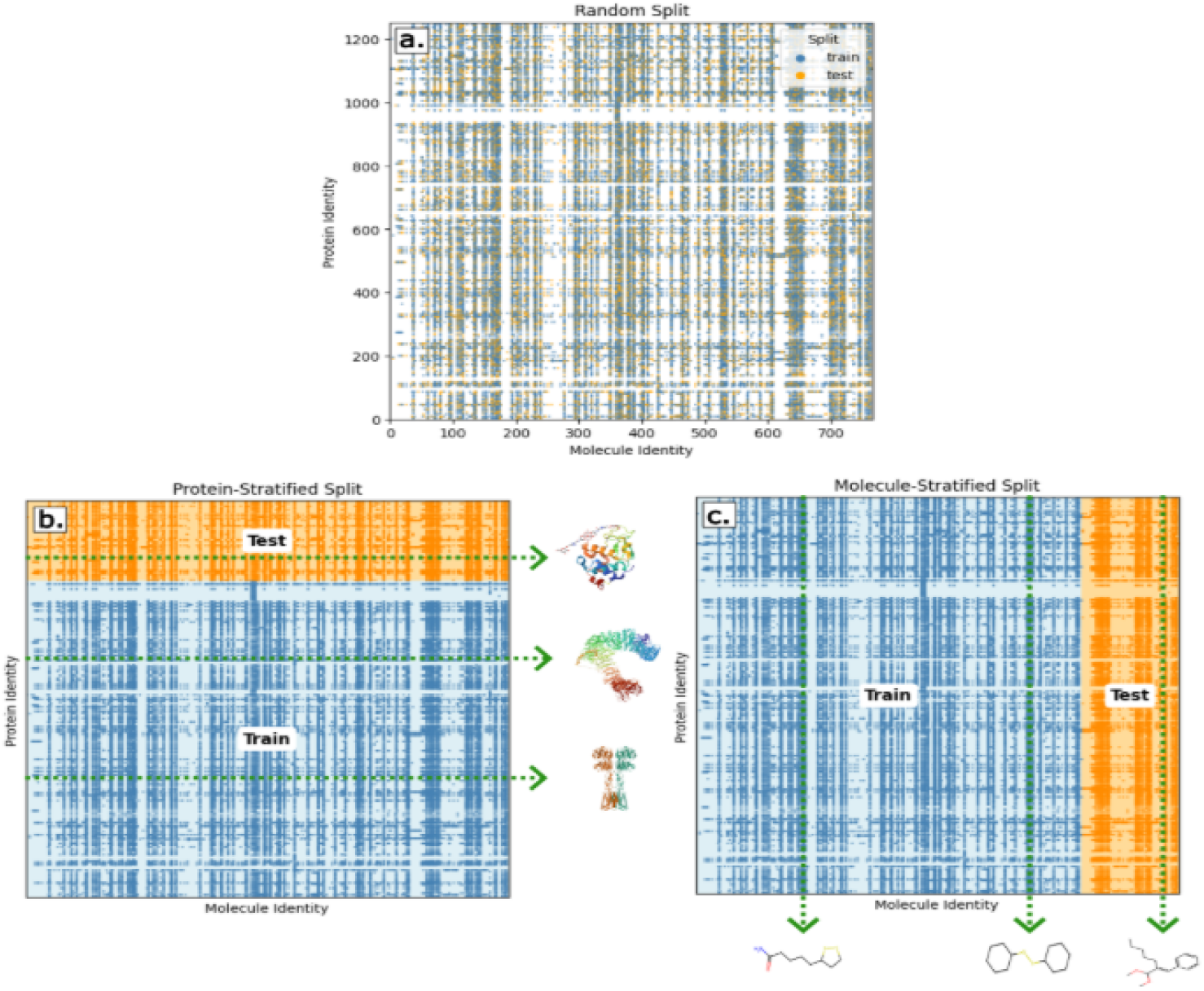
Data stratification strategies. Each VOC identity in a VOC-OR is plotted against its OR pair. A. The random splitting shows the train and test set contain data points that share VOC or OR identities. B. The dataset is stratified by the OR identity, meaning no ORs found in the train set are present in the test set. C. The dataset is stratified by the VOC identity, meaning no VOCs found in the train set are present in the test set

A two-layer fully connected neural network was used to classify pair binding. The model was optimized using Binary Cross Entropy (BCE) loss with logits, with a varying positive class weight hyperparameter to adjust for large class imbalance in M2OR. Area Under the Curve of the Receiver

Operating Characteristic curve (AUCROC) was the primary metric to appraise the model’s ability to discriminate binding pairs, especially in a class-imbalanced context. The data was split using a 5 fold cross validation split, giving a 20% dataset test size with 15% of the resulting train set being used for validation.

In addition to training on BCE, a neural network was developed using a contrastive loss function. The former is referred to as the “BCE” model and the latter is the “contrastive” model. Contrastive training requires that each batch given to the machine learning model contains at least one positive pair.

For each contrastive training batch, a stochastic sampler selected a random VOC-OR identity pair to act as an “anchor”. The sampler then selected additional samples that shared either a protein or VOC with the anchor and labeled them as positive samples. Negative samples were samples that did not share either a VOC or an OR with the anchor pair. A full epoch using the sampler was defined after all OR or VOC identities – depending on the choice of OR or VOC stratification – in the dataset had been sampled once. Throughout training, the sampling probability of selecting identities was altered: a linear increase in probability of selecting samples that were not selected, including an exponential term for identities not selected that epoch, and a geometric decay of the identities chosen. The linear increase promotes undersampled points, the geometric decay discourages sampling the same identities many times in a small amount of batches, and the exponential term encourages full coverage of identities. A robust explanation can be found in Supplementary Information.

The contrastive model used the latent space to create the contrastive loss criterion. A triplet margin loss function was applied to the latent space in the hidden layer of the network. In contrastive experiments, the same train/test/val split ratios were used and hyperparameters for the sampler weighting scheme and the margin and weight of the triplet loss were added to the hyper-parameter study.

Each experiment was comprised of an encoder pair (ProtBERT or Integers for the protein, ChemBERTa or Morgan fingerprints for the VOC) and a data stratification strategy (Random, Protein-stratified, or VOC-stratified), with twelve total experiments to explore the combined space. For each experiment, the model’s hyperparameters were tuned on a single random seed using Optuna. BCE model experiments used 100 trials and were allowed to train up to 100 epochs, with early stopping using patience of 10 epochs and a tolerance of 0.01 measured using the validation AU-CROC. Contrastive model experiments used 10 trials and were allowed 10 epochs, with a patience of 15 steps, where every step comprises 150 batches, validated in the same way. This discrepancy was due to the larger computational complexity, and the fast convergence of hyperparameters observed in initial contrastive experiments.

After optimal hyperparameters were selected, the tolerance term and patience were removed, and the model was evaluated on the test set. The entire training and evaluation regimen was then repeated with 5 random seeds, with 5 fold cross validation - five results are reported for each experimental condition

## 4 Results

For a neural network classifying binding affinities in iOBPdb with no contrastive loss, the encoder methods were the primary predictors of performance, independent of the data splitting method. Figure 2 summarizes the results of random seed trials. Models trained with the Morgan fingerprint and AA Integer encoders performed the best with an AUCROC of 0.91 for all splits. This was followed by the ChemBERTa-AA Integer encoder, then the ChemBERTa-ProtBERT, then the Morgan fingerprint-ProtBERT encoder models with 0.9, 0.87, and 0.84 AUCROC respectively. Transformer encoders had diminished performance on iOBPdb.

**Figure 2.**
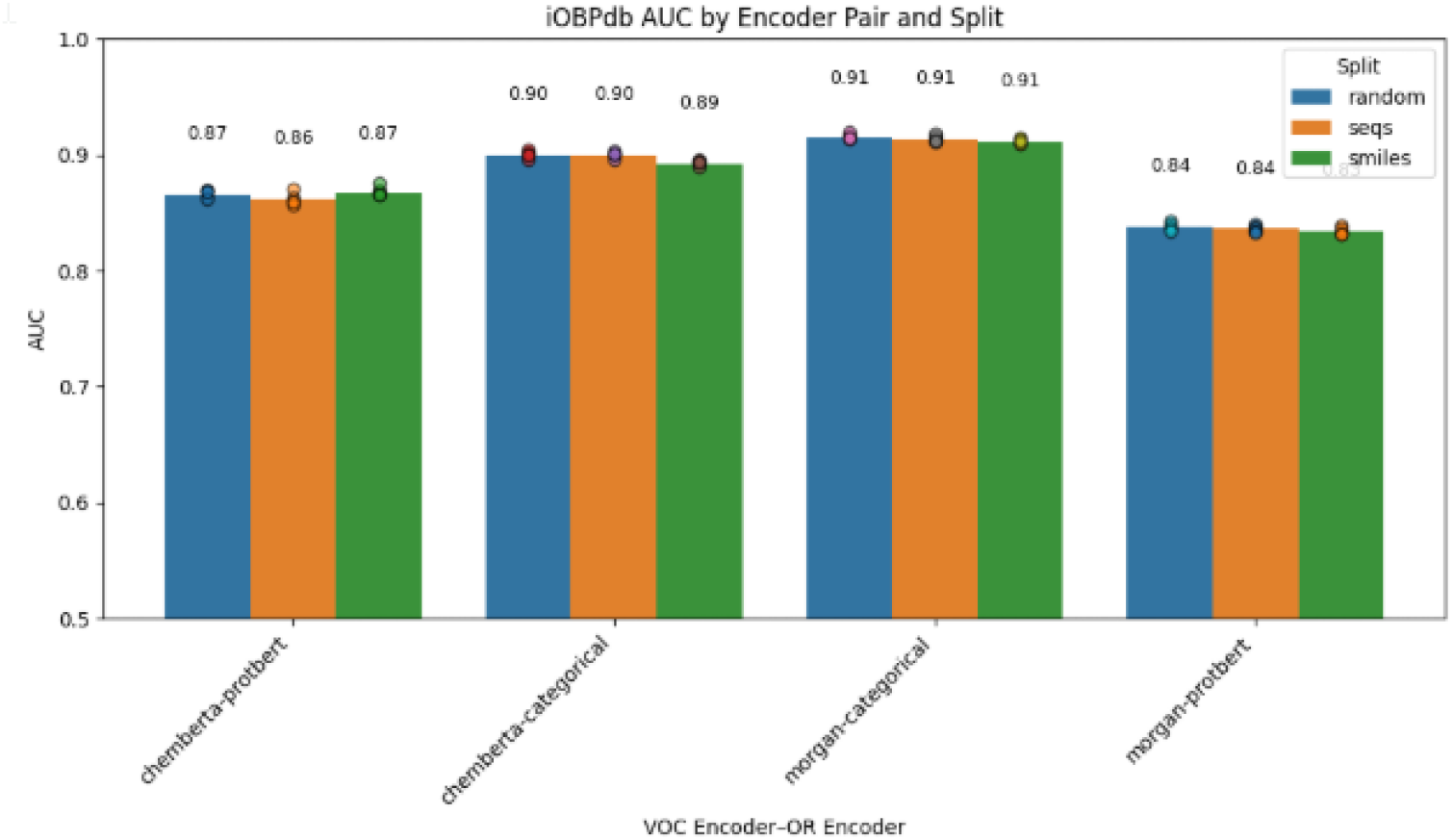
AUCROC across iOBPdb experiments over 5 random seed trials. Transformer-based encoders did not yield better performance.

For the neural network on M2OR with no contrastive loss, data splitting strategy had a major effect on performance, interdependent with encoder choice. Figure 3 summarizes the results of random seed trials. Models trained on randomly split data outperformed models trained on VOC and OR stratified splits by a 0.1 to 0.2 point margin. The Morgan-Categorical model under a random split performed the best with an AUCROC of 0.88. Under an OR-stratified split, the Morgan-ProtBERT pair model performed the best with an AUCROC of 0.81. For the VOC-stratified split, the ChemBERTa-Categorical pair model performed the best with an AUCROC of 0.78.

**Figure 3.**
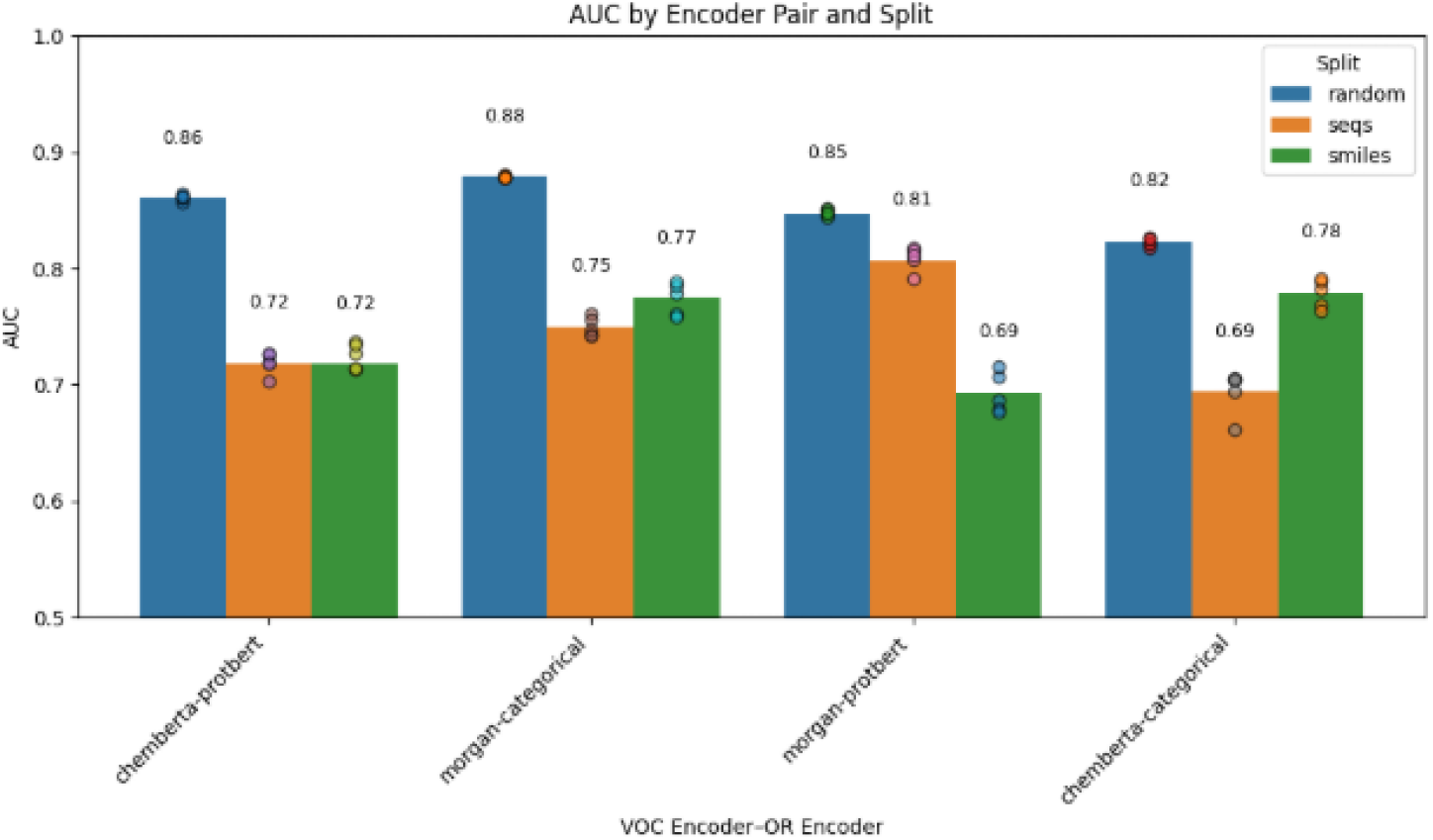
AUCROC across M2OR experiments over 5 random seed trials. Random splitting performed the greatest overall, indicating the model’s reliance on statistical overlap of the train and test set. Performance on the VOC and OR stratified splits favors pretrained transformers that have larger representation capacity along the axis of stratification.

Figure 4 shows the evolution of the validation set AUCROC throughout the training process for all experiments. Models using the random split converged quickly, reaching stable AUCROC within 10-15 epochs. By comparison, models trained using the VOC- and OR-stratified splits showed slower convergence, larger variance across random seeds and cross validation folds, with a lower AUCROC.

**Figure 4.**
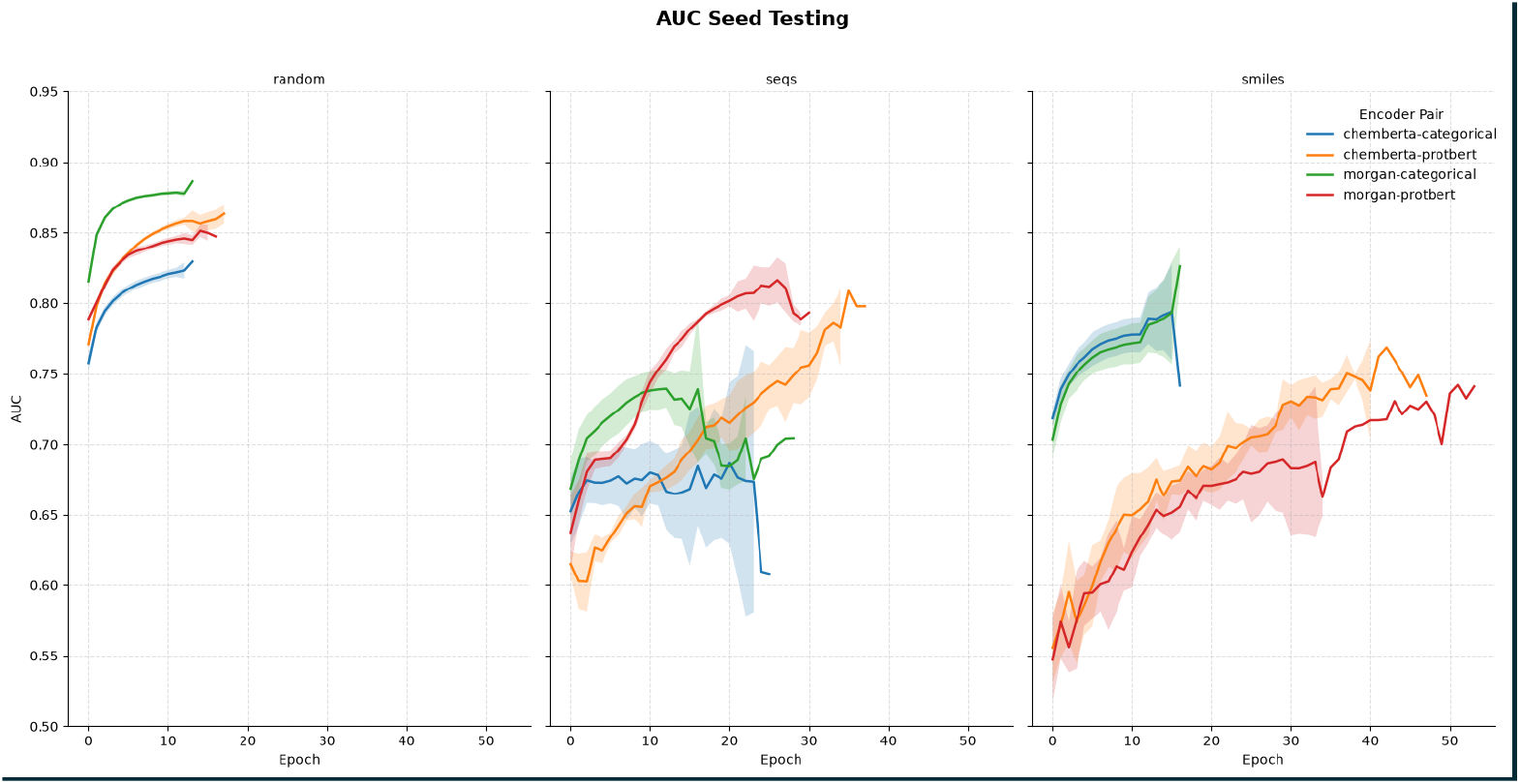
AUCROC over epochs for all experiments. Models using randomly stratified splits had stable, quick convergence, while the models VOC and the OR splits had more chaotic convergence, with longer tails.

For the contrastive experiments, the dependence on data-stratification largely disappeared. Using the Morgan-Categorical encoder pair and the ChemBERTa-ProtBERT encoder pair performed the best with 0.88 AUCROC score while the other encoders performed at 0.83 AUCROC score. These values remained relatively constant across random, VOC-, and OR-stratified splits. Results from this are summarized in a bar chart in Figure 5.

**Figure 5.**
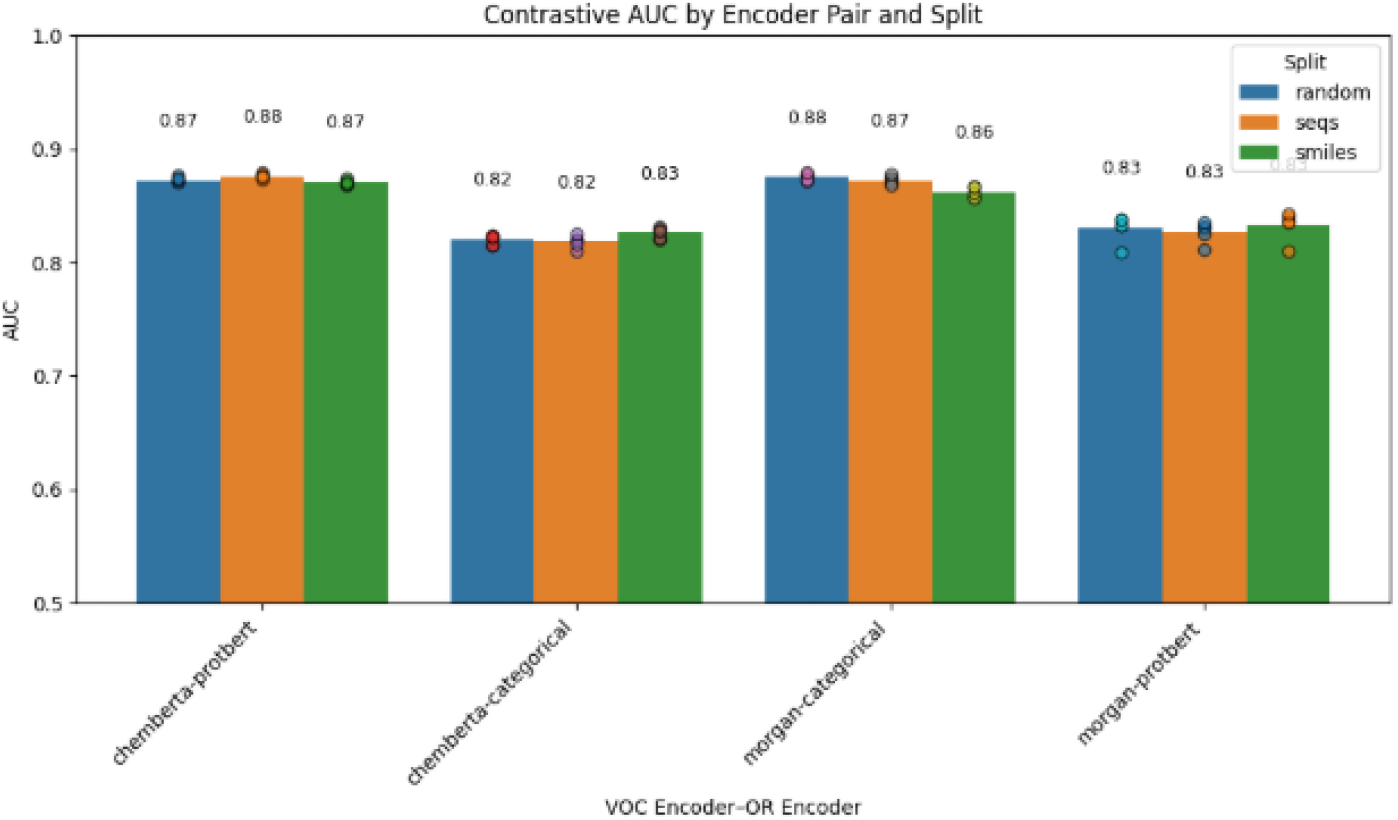
AUCROC across contrastive experiments. Model performance does not significantly vary with on stratification strategy .

Figure 6 shows the AUC training curves throughout the contrastive training process. Contrastive models displayed more consistent convergence and lower inter-seed and inter-fold variance. Convergence was generally slower; considering the difference of steps to epochs, the contrastive method processed many more samples than the naive method.

**Figure 6.**
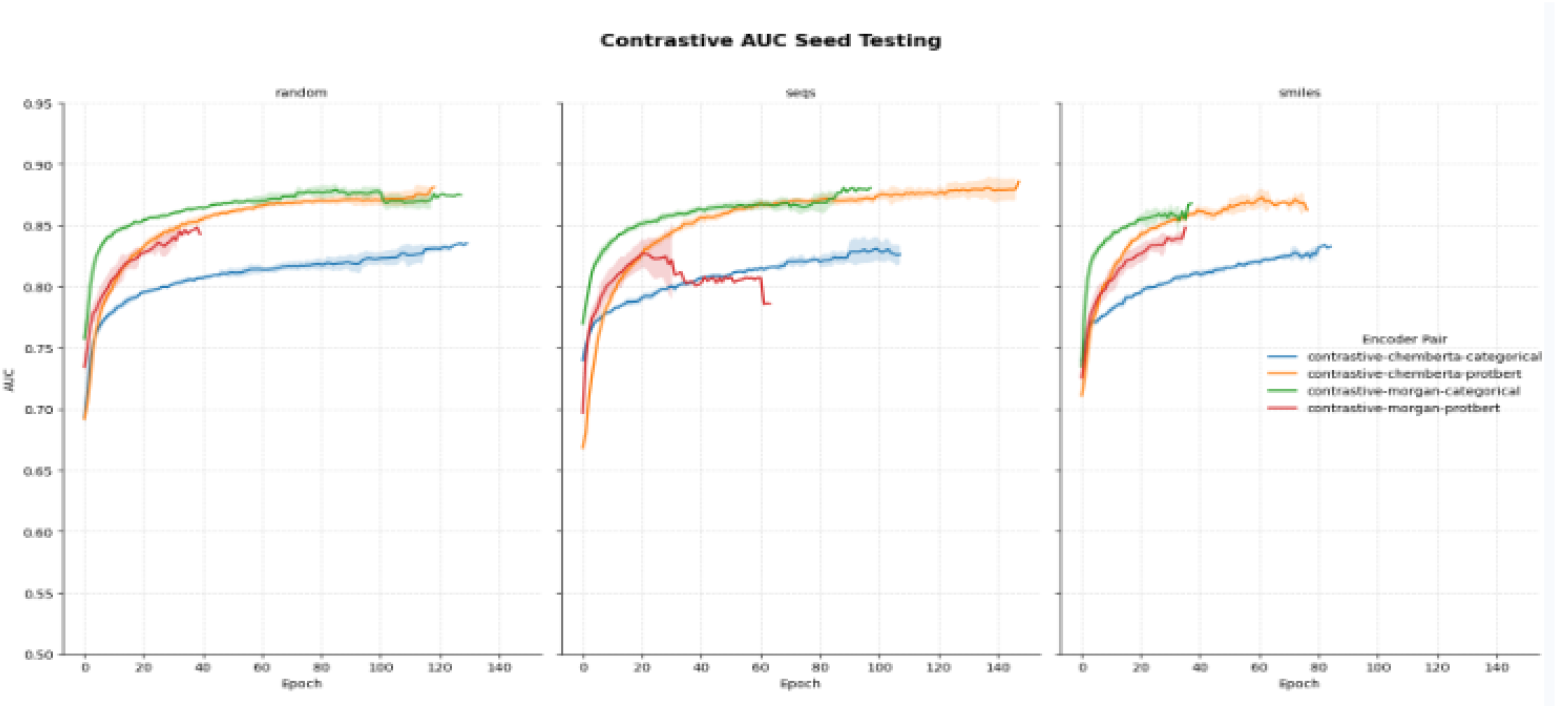
Mean AUC of contrastive models over 5 folds. The shaded region shows the standard deviation of all folds at that epoch. All splitting strategies tightly converge over training.

## 5 Discussion

This study observed the impact of data partitioning, encoding strategies, and contrastive vs. traditional loss functions on binding affinity prediction between proteins and VOCs in two datasets. M2OR is a class-imbalanced dataset with a positive percentage of 6% and contains considerable label noise from unifying many heterogeneous biological assays, which may force the model to find an optimization path dominated by dataset-driven effects and set an upper limit of model performance. [19, 7] Chithrananda et al. report a state of the art validation AUCROC on M2OR of 0.8897, using a GCN to encode the VOC and a protein language model (ESM-2) to encode the OR with a learned attention pooling block and a neural network classifier (MolOR)[23]. McConachie et al. build on this work by implementing fine tuning through low rank adaptation and report that ProSmith can achieve an AUCROC of 90.47 on M2OR [24]. However, both only evaluate the models on the random split, and the ProSmith AUC was reported on a variant of the M2OR dataset that removed VOCs and ORs with low representation. Across reports, performance in AUCROC remains near or below 0.9.

The BCE model’s AUC of 0.88 on randomly split data approached state-of-the-art performance with comparatively simple encoders and models, but performance sharply diminished when using VOC- or OR-stratified splits. With a random split, the algorithmic encoder pair performed the best while other encoding methods trailed by a small margin. Encoder pair performance was also affected by data stratification: models with randomly stratified data favored low representational capacity, but when specific proteins or VOCs were held out from the training set, encoding methods with more representational capacity along the stratification axis were favored. Maximum AUCROC of 0.88 was achieved on the model trained on the randomly split data, with an AUCROC of 0.81, and 0.78 for neural network models with the OR- and VOC-stratified data respectively suggest that lack of biochemical features diminishes the model’s capability to generalize. This result indicates that model choice should address the demands of the test task, with increased representational power needed to extrapolate to novel molecules along the protein or VOC axis. In this model-limited regime, we posit the model may rely primarily on statistical correlation of features in the training and evaluation sets, rather than a reliance on chemical and biological molecular properties. Sensitivity to encoding method highlighted a key trade off in biomolecular transfer learning.

The algorithmic encoding methods (Morgan, categorical) contain no bias; they simply convert the inputs to a naive representation based on the algorithm provided. Meanwhile, the transformer based representations (ProtBERT, ChemBERTa) import a manifold with rich representation capacity, but they inherit the bias from their training dataset. Ultimately what was observed was that the transformer encodings were less flexible than the bias-free algorithmic encodings, and when presented with tasks to maximize the classification of motif overlap in the random split, performed marginally worse. When these encodings were used on the stratified tasks, the representation power for the stratified feature and the flexibility for the other feature yielded much better performance. It’s clear that the bias of the transformer encodings make them unsuitable as a blanket solution for these biochemical tasks, and can even be a boon due to their strong existing bias to support prediction on inputs from outside the dataset.

When a contrastive objective was introduced to the model, performance became insensitive to the dataset splitting method. Unlike the BCE model, which struggled when it could not rely on memorizing motifs from the training distribution, models trained with a contrastive criterion showed apparently no such degradation. Performance was still bounded by a 0.88 limit; the model may have improved its ability to generalize, but the performance ceiling remained intact.

This work has shown that many model configurations can achieve 0.88 AUCROC, but their ability to generalize to novel binder systems is highly variable. Implementing powerful pretrained encoders can partially improve generalization for the corresponding molecular species. However, introducing a contrastive loss term to the model provided robust improvement on unseen species across all encoder pairs. One can see a single data point as modeling not the pair of VOC and OR, but representing the shared physicochemical binding interface between them. The objective in M2OR presents a challenge in building a compatibility manifold that well represents this binding topology using only semantic labels. Contrastive learning may provide a strong inductive bias for this task. Manifolds that likely express the same physicochemical binding topology motifs from either the VOC or the OR as augmentations for contrastive regularization directly supports alignment of a compatibility manifold through these augmentation labels. The structured latent representation created by the contrastive training yielded generalizable representations to both unseen protein and molecular physicochemical motifs, can match state of the art performance with models with much less representation power.

This study establishes a clear correlation between performance, data stratification, and architecture design; future work may seek to further validate and effectively apply these insights to olfactory prediction in general. The lack of data split sensitivity in iOBPdb models suggests that M2OR either represents a more difficult to model or underrepresented class of binding events. With the rise of computational bio molecular datasets, data availability is steadily increasing and opportunity exists to explore these conclusions on datasets that are more robust. Future work can use embedding similarity analysis to better interpret features and may open the door to self-supervised methods that generate better features pertinent to both biological binding and machine learning performance. Effort can be taken to analyze which proteomic or chemical motifs drive the difficulty in binding prediction, which may elucidate the motifs that are underrepresented in currently available data. By doing so, data collection procedures can be optimized, enabling synergistic development of data and model facets of VOC-OR binding.

## 6 Conclusion

In the M2OR olfactory receptor binding dataset, distributional overlap of the train and test set has a significant effect on VOC-OR binding prediction performance compared to specific choices of encoder and model regularization. Through systemic isolation of effects from data partitioning, data sampling, encoder choice, and model architecture, we show that model performance in VOC-OR binding prediction on M2OR is degraded when data partitioning forces the model to predict novel proteins or VOCs. In this regime, using transformer based encoders for the novel prediction feature encoder, with a flexible algorithmic encoder yielded the best results. A contrastive augmentation labeling scheme was developed by defining positive pairs through shared molecular constituents. By introducing a contrastive objective criterion to the loss function, the contrastive model was able to perform well across VOC-stratified, OR-stratified, and random splits, regardless of encoder choices. These findings emphasize the importance of dataset biases and model evaluation practices for developments in model design to effectively translate architectural advancements to engineering insights and implementation. This study contributes nuance to model performance interpretation on two VOC-OR datasets, showing that machine learning development is highly contextualized by the choice of data splitting method, architecture and regularization techniques. The contrastive learning method, which matched state of the art performance, is a promising technique to stabilize model training and reduce sensitivity to specific data split choices, and it hints at an underlying relational structure for olfactory pairwise interaction datasets that may be relevant to other binder-ligand machine learning tasks.

