## Supplementary Information for "Contrastive Learning Improves Generalization on M2OR Olfactory Binding Dataset"

### Supplementary Information: Identity-Aware Contrastive Sampling

#### 1 Overview

Contrastive learning requires each batch of samples to contain an anchor with a controlled set of same-class ("positive") and different-class ("negative") examples. In this case, positives are defined as samples that share a constituent binder and the same binding classification, and negatives are any sample that does not meet either one of those criteria. These positive samples may have similar binding motifs that contribute to its binding affinity, allowing the contrastive criterion to help cluster different motifs from different molecular constituents. For the construction of the anchor from volatile-organic-compound (VOC) ligand and olfactory-receptor (OR) pairs, it may take a long time to select a sample from a uniform distribution. To reduce the amount of time while respecting the frequency structure of the dataset, we developed an identity-aware sampler that for each batch it selects an anchor from an adaptively reweighting distribution and builds a batch with specific amounts of positive, negative, and hard negative samples around the selected anchor.

#### 2 Sampling Algorithm

##### 2.1 Notation

Let $\mathcal{D} = \{(v_i, o_i, y_i)\}_{i=1}^{N}$ denote a training dataset of $N$ samples, where $v_i$ is a VOC identity, $o_i$ is an OR identity, and $y_i \in \{0, 1\}$ is their binding classification. Let$V = \{v_i\}$ and $O = \{o_i\}$ denote sets of unique VOC and OR identities in $\mathcal{D}$.

##### 2.2 Batch construction

For a selected anchor example $a = (v_a, o_a, y_a)$, we define

$$\mathrm{Local}(a) = \{\, i : v_i = v_a \;\vee\; o_i = o_a \,\} \tag{1}$$

$$\mathrm{Positive}(a) = \{\, i \in \mathrm{Local}(a) : y_i = y_a,\; i \neq a \,\} \tag{2}$$

$$\mathrm{Negative}_{\mathrm{local}}(a) = \{\, i \in \mathrm{Local}(a) : y_i \neq y_a \,\} \tag{3}$$

$$\mathrm{Negative}_{\mathrm{global}}(a) = \{\, i \notin \mathrm{Local}(a) : y_i \neq y_a \,\} \tag{4}$$

At least `min_pos` positives and `min_neg` negatives are guaranteed for every anchor by directly sampling $\mathrm{Positive}(a)$ and $\mathrm{Negative}_{\mathrm{local}}(a) \cup \mathrm{Negative}_{\mathrm{global}}(a)$. The remainder of the batch is filled from a combination of samples from $\mathrm{Local}(a)$ and $\mathrm{Negative}_{\mathrm{global}}(a)$, with the composition of local and global samples being controlled by the `global_ratio` hyperparameter.

#### 2.3 Adaptive identity reweighting

Each VOC identity $v$ and OR identity $o$ is initially assigned a weight, $w_v = w_o = 1$. An anchor example $i = (v_i, o_i)$ is drawn with probability of the summed weight of its two constituent identities:

$$P(i) \;=\; \frac{w_{v_i} + w_{o_i}}{\sum_{j=1}^{N} \left(w_{v_j} + w_{o_j}\right)}. \tag{5}$$

Following the selection of anchor identities, every identity's weight, $v$, is updated in the following way for the selected anchor, $v^*$:

$$w_v \;\leftarrow\; \begin{cases} \gamma\, w_v & v = v^* \\ w_v + \dfrac{\alpha}{N} + \alpha\left(1 - e^{-4c/N}\right) & v \neq v^*,\; v \notin \mathcal{C}_V \\ w_v + \dfrac{\alpha}{N} & v \neq v^*,\; v \in \mathcal{C}_V \end{cases} \tag{6}$$

where $\mathcal{C}_V$ is the set of VOC identities already selected as an anchor at least once during the current epoch, $c = |\mathcal{C}_V \cup \{v^*\}|$ is the number of unique VOC identities covered so far (including current), $\alpha \in (0, 1)$ is a linear growth rate hyperparameter, and $\gamma \in (0, 1)$ is a geometric decay-rate hyperparameter. $N$ is the total number of training examples, so the per-step growth increment and the coverage ratio driving the exponential term are both scaled relative to dataset size to tie the reweighting schedule to the overall size of $\mathcal{D}$.

Three distinct behaviors are encoded by Eq. 6. An identity's weight decays geometrically by a factor of $\gamma$ each time it is selected to disincentivize repeated sampling of high representation identities. An identity that has not been selected during that epoch receives an additional exponential boost term that grows as the fraction of the manifold already covered increases, creating accelerating pressure to sample previously unseen identities as an epoch progresses. An identity that meets both of the previous criteria is only linearly incremented to promote diversity while still respecting frequency structure. Because decay is applied once per selection event, frequently selected identities are suppressed in proportion to how often they are actually drawn and rare identities that are eventually selected receive only a small multiplicative penalty while high representation identities are suppressed.

One epoch of sampling continues until every unique VOC and OR identity has been selected as an anchor at least once.

Table 1: Sampler configuration used to generate the validation results shown in Fig S1-S3. In training these hyperparameters are selected with hyperparameter search.

| Hyperparameter | Value |
|---|---|
| Batch size | 64 |
| Minimum positives per anchor | 2 |
| Minimum negatives per anchor | 2 |
| Linear growth rate ($\alpha$) | 0.05 |
| Geometric decay rate ($\gamma$) | 0.7 |
| Global negative ratio | 0.3 |

#### 3 Baseline Validations

To isolate the effect of this sampling method from the batch construction, we compared the full selection mechanism described above – termed "weighted" – against a baseline in which every identity's sampling weight is held fixed at 1 for the duration of the run – termed "naive", equivalent to drawing anchor examples uniformly at random over $\mathcal{D}$ with no reweighting. Both conditions were run on identical real ligand-receptor identity and binding-label data drawn from the training split of the M2OR dataset, with the same random seed and number of selection steps, so that any difference in outcome may be ascribed to the reweighting mechanism.

##### 3.1 Results

Figure S1 shows: sampling weights for two high-frequency and two low-frequency VOC identities over steps (a), cumulative identity coverage over steps (b), per-anchor positive and negative sample count (c), and a comparison of the frequency of identities and how many times they were picked as an anchor(d). OR identity coverage approached saturation within the sampled epoch, while VOC coverage had not yet fully plateaued over the same number of selection steps.

Figure S2 compares mean anchor-selection counts per identity, binned by row count, between the naive and weighted conditions. In the naive condition, selection count increases directly with row count, as expected from uniform row sampling. Under the weighted condition, identities in the highest row-count bins are selected less often on average than under naive sampling, while lower-frequency identities are selected much more often, reducing the dependence of selection frequency on raw dataset representation.

Figure S3 compares identity coverage between the two conditions over an equal number of steps. The adaptive method reaches a given level of identity coverage earlier than the naive method. The shape of the cumulative distribution functions plotted are similar, indicating they act similarly, only the adaptive method saturates less. This promotes diversity as intended, as unseen or sparsely sampled samples are seen more often by the model. Due to the fact that altering the frequency structure could impact the model's performance, the sampler reweighting was tuned as a hyperparameter.

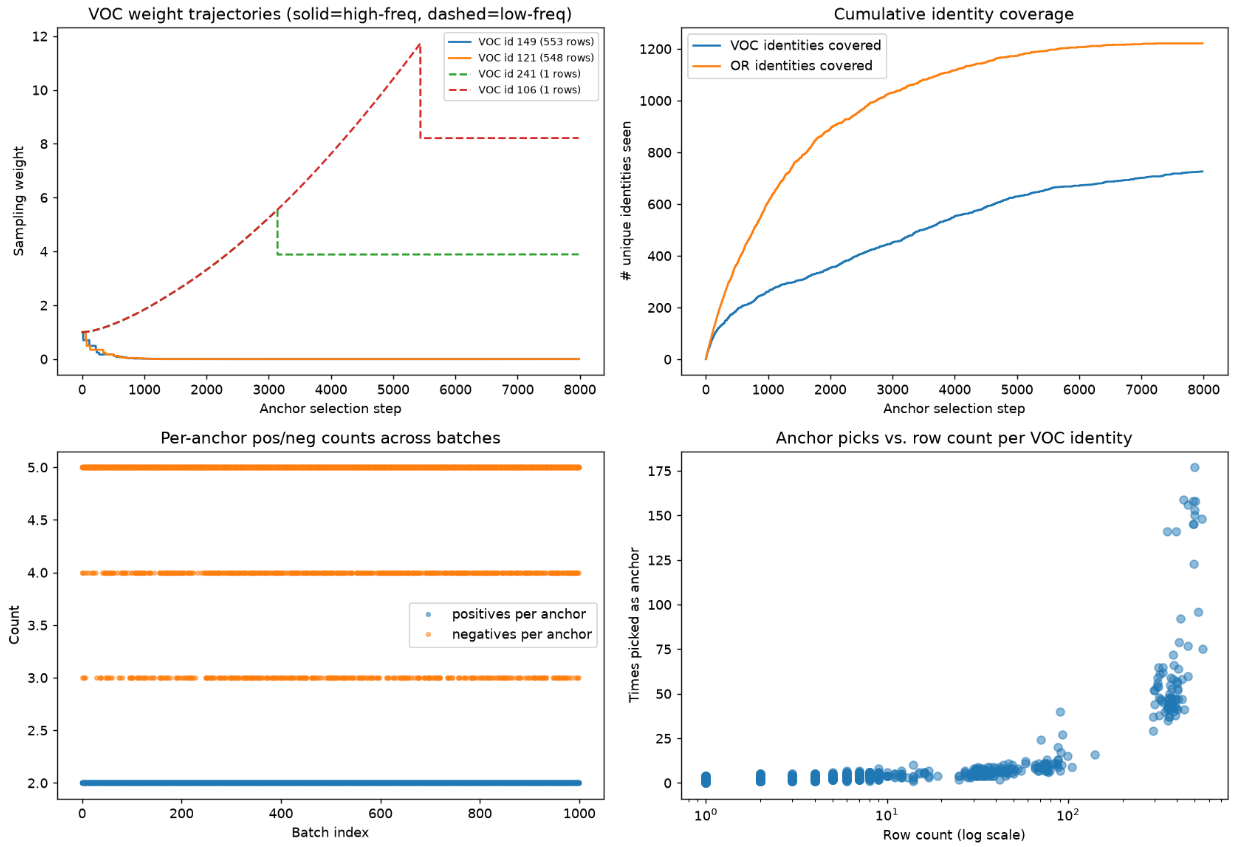


Figure 1: Diagnostics from one full epoch of the weighted sampler on real training data. (a) Sampler weight trajectories over the course of the epoch for two high-frequency and two low-frequency VOC identities, illustrating the geometric decay applied on selection and the accelerating growth applied to not-yet-selected identities. (b) Cumulative number of unique VOC and OR identities covered as a function of selection step. (c) Positive and negative examples per batch across steps to confirm desired batch structure guarantees. (d) Total number of times each VOC identity was selected as an anchor over the full epoch, plotted against that identity's row count in the training set in log scale.

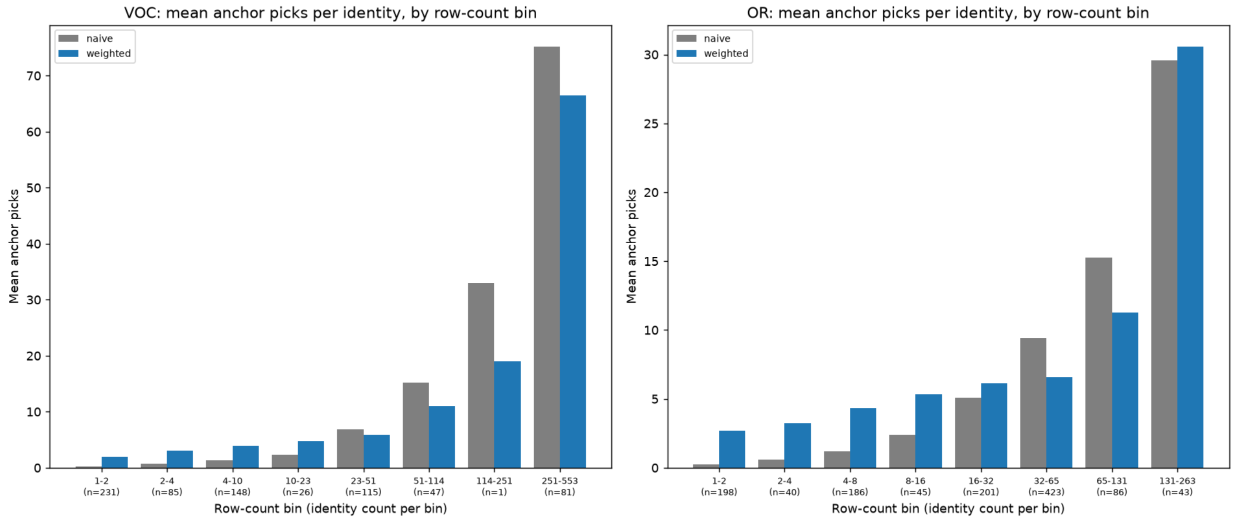


Figure 2: Mean number of anchor selections per identity, binned by log-space row-count ranges, comparing naive (uniform row sampling, no reweighting) and weighted sampling. Left: VOC identities. Right: OR identities. Bin labels indicate the count range of the bin and the number of unique identities that fall in that bin ($n$). Weighted sampling reduces the dependence of selection frequency on dataset representation relative to naive sampling.

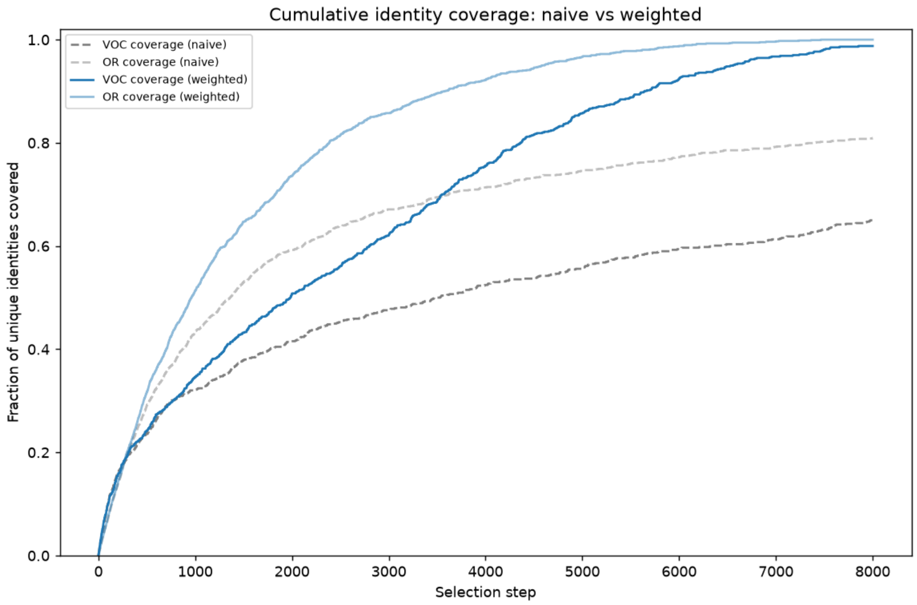


Figure 3: Cumulative fraction of unique VOC and OR identities selected as an anchor at least once, as a function of selection step, comparing naive and weighted sampling on identical training data and random seed. Weighted sampling reaches a given level of identity coverage earlier and experiences less saturation than naive sampling for both identity types throughout the sampled range.
